# Comparative transcriptomic analysis of whole blood mycobacterial growth assays and tuberculosis patients’ blood RNA profiles

**DOI:** 10.1101/2022.06.24.497240

**Authors:** Petra Bachanová, Ashleigh Cheyne, Claire Broderick, Sandra M. Newton, Michael Levin, Myrsini Kaforou

**Affiliations:** Department of Infectious Disease, Faculty of Medicine, Imperial College London, Norfolk Place, London W2 1PG; MRC Centre for Molecular Bacteriology and Infection, Department of Life Sciences, Imperial College London, Faculty of Natural Sciences, South Kensington Campus, London SW7 2AZ; Centre for Paediatrics and Child Health, Imperial College London, Faculty of Medicine, South Kensington Campus, London SW7 2AZ

## Abstract

*In vitro* whole blood infection models are used for elucidating the immune response to *Mycobacterium tuberculosis* (*Mtb*). They exhibit commonalities but also differences, to the *in vivo* blood transcriptional response during natural human *Mtb* disease. Here, we present a description of concordant and discordant components of the immune response in blood, quantified through transcriptional profiling in an *in vitro* whole blood infection model compared to whole blood from patients with tuberculosis disease. We identified concordantly and discordantly expressed gene modules and performed *in silico* cell deconvolution. A high degree of concordance of gene expression between both adult and paediatric *in vivo-in vitro* tuberculosis infection was identified. Concordance in paediatric *in vivo* vs *in vitro* comparison is largely characterised by immune suppression, while in adults the comparison is marked by concordant immune activation, particularly that of inflammation, chemokine, and interferon signalling. Discordance between *in vitro* and *in vivo* increases over time and is driven by T-cell regulation and monocyte-related gene expression, likely due to apoptotic depletion of monocytes and increasing relative fraction of longer-lived cell types, such as T and B cells. Our approach facilitates a more informed use of the whole blood *in vitro* model, while also accounting for its limitations.

## Introduction

According to the World Health organisation (WHO) Global Tuberculosis Report 2021^1^, there was an estimated 1.3 million deaths caused by tuberculosis (TB) in 2020. Despite being curable and preventable, TB has lingered at the top of the killer communicable disease list for many decades. Limited access to healthcare in poorer countries, an absence of diagnostic tests with sufficient sensitivity, robustness and affordability, and the lack of a universal vaccine capable of conferring immunity to both adults and children are contributing to the strain. Progress in the latter two is hindered by the elusive host immune response to TB, and the incomplete understanding of immunological mechanisms modulating an individual’s ability to fight the infection.

TB is caused by the airborne pathogen, *Mycobacterium tuberculosis* (*Mtb*). Upon inhalation *Mtb* is faced with the first line of defence in the form of “professional” phagocytes (macrophages, neutrophils, and dendritic cells). If successful, *Mtb* infects these cells and rapidly proliferates within them. Once TB infection is established it may stay dormant for years in a delicate interplay between the host immune system and the bacilli with one quarter of the world’s population^2,3^ estimated to be infected with *Mtb*. The individual is at highest risk of developing TB disease within the first two years after infection but can remain at risk for their lifetime.^4^ Younger children are more likely to progress to primary infection. In older children and adults, progression from latent infection to TB disease may occur as a consequence of a weakened host immune response, through factors such as co-infections, other diseases or ageing, as well as pathogen immune-escaping mechanisms. These include evading degradation within the phagolysosomes, delaying the activation of the adaptive immune response, and mutating its surface antigens to evade T-cell recognition^5–7^.

Whole blood transcriptomics can improve our understanding of the host immune response to *Mtb* infection and progression to disease. Blood gene expression profiling of patients with TB has highlighted *Mtb* specific transcriptional changes associated with cellular pathways involving Interferon gamma (IFN-γ) and T cell receptor signalling and proliferation^8^. As the patterns of expression in the transcriptional profiles are specific to different pathogens, the last decade has focussed much attention to exploring the role of gene expression signatures in disease diagnosis and prognosis in paediatric and adult patient cohorts with pulmonary TB or extrapulmonary TB ^9–12^. Gene signatures identified between active TB and other disease control cohorts are currently being investigated for use as biomarkers in order to form the basis of more sensitive diagnostics tests^13^.

*In vitro* infection models provide a highly controlled way of studying host-pathogen interaction and allow for robust, reproducible, and translatable research. Infection models have played an important role in current understanding, treatment, and prevention of many infectious diseases, including TB disease^14–17^. In comparison to studying natural infection in patient cohorts they can be less confounded by various factors including pathogen strain and dose, can be broader in scope, cost-efficient, less time-consuming, and can provide longitudinal measurements. However, the relevance of any given model to the human infection and disease needs to be evaluated, as there is the risk that it may not recapitulate biological processes underpinning natural infection. Understanding how the *in vitro* model resembles *in vivo* infection is essential for extrapolating the model’s experimental findings to *in vivo* natural infection and inform vaccine development and host-directed therapies.

Whole blood infection assays (WBA) are a reliable *in vitro* method used to assess human cellular immune responses to *Mtb* or *Mtb* antigens. They have been shown to better model the diverse cellular interactions that manifest during the immune response to infection compared to peripheral blood mononuclear cells (PBMC), with higher cell viability and encompassing all immune cell types and non-cellular components of the human blood^17^. WBAs have also been used to evaluate *Mtb*-specific T cell responses, their breadth and specificity^15^. More recently, bulk human RNA transcriptional profiles from an *in vitro Mtb* infection WBA were employed to assess longitudinal host responses to *Mtb*, revealing a widespread immune suppression over time^6^. Whilst this model provided important new data in the study of the host immune response to *Mtb* with implications for the discovery of new vaccines and therapeutics, its similarity to natural host *Mtb* infection has not been previously assessed.

Recently a comparative transcriptomics approach for human and murine transcriptional responses to *Mtb* infection identified the drivers of concordance and discordance between the two systems^14^. This method evaluates the degree of similarity based on the direction of gene regulation (up or down) weighted by the magnitude of the effect size and associated significance.

Here, we aim to identify the similarities and differences at the transcriptional level of an *Mtb* infection WBA and natural *in vivo* host response in adults and children with TB disease. Identifying biological pathways that drive concordance and discordance in these datasets is of great importance to facilitate a better understanding of the *in vitro* WBA system, and the components of the *in vivo* infection that are recapitulated over time. Lastly, we employ *in silico* cell deconvolution prediction to elucidate the role of possible differences in immune cell populations in driving concordance and discordance between the *in vivo* and *in vitro* systems.

## Methods

Data analysis was performed in R version R-4.0.3^18^, and a script including all analytical steps is available upon request.

### Data Acquisition

Previously published whole blood gene expression microarray datasets were downloaded from Gene Expression Omnibus database (accession number for *in vitro* dataset: GSE108363^6^, for adult *in vivo* dataset: GSE37250^19^, for paediatric *in vivo* dataset: GSE39941^10^). The gene expression microarray datasets had been generated using Illumina HumanHT-12 v4 Expression BeadChip microarrays.

The *in vitro* dataset derives from a study of healthy adult donor peripheral whole blood infected with a bioluminescent strain of *Mtb* (H37Rv *Mtb lux*) using a WBA. This assay incubates infected and uninfected whole blood over a time-course allowing for analyses, including bulk gene expression microarray analysis in this instance, to be performed at different time-points. In this dataset whole blood samples included 44 infected and 52 uninfected controls at 6 h, 24 h, 48 h, 72 h, and 96 h in the discovery and validation cohorts (n=4 and 6 respectively).

*In vivo* datasets were probed for gene expression changes in adults and children diagnosed with active TB, relative to LTBI. The paediatric dataset consisted of samples from patients from South African, Kenyan, and Malawian cohorts, with 82 active TB samples and 57 LTBI samples (mean age 5.1 years, median age 3.3 years, min = 3 months, max = 15.4 years). The adult dataset consisted of a South African and Malawian cohort, with 97 active TB samples and 82 LTBI samples (mean age 34.8 years, median age 29.7 years, min = 17.8 years, max = 78.9 years).

### Data pre-processing

The datasets were quantile normalised across arrays and transformed to a logarithmic scale (base 2). Batch correction was performed using ComBat function from the sva package^20^.The *in vivo* data sets were corrected for the study sites and the *in vitro* dataset was corrected for discovery and validation cohorts.

For *in vivo* datasets, only the samples from Human Immunodeficiency Virus (HIV)-uninfected participants with culture-confirmed TB were included in the analysis. LTBI individuals were used as a proxy for controls. Sample GSM914578 from the adult dataset (GSE39941) was excluded due to missing data. For the *in vitro* dataset, only *Mtb*-infected and uninfected samples were used. Principal component analysis was performed using the PCA tools package^21^. Other packages used for data processing included: pheatmap^22^, ggplot2^23^, ggrepel^24^, magrittr^25^, RColorBrewer^26^, mgsub^27^, data.table^28^, biomaRt^29^, taRifx^30^, reshape^31^, edgeR^32^.

### Differential gene expression analysis and pathway analysis

Limma R package^33^ was used for differential gene expression analysis. The linear models accounted for disease group, study site, sex, and age, in the *in vivo* datasets and for treatment condition (*Mtb*-infected and uninfected whole blood) and treatment time in the *in vitro* dataset. Gene mapping was performed using biomaRt^29^ and differentially expressed genes were visualised using the Enhanced Volcano package^34^. Log_2_ fold change (FC) threshold was set at 0.5 and – 0.5 -and Benjamini-Hochberg adjusted p-value was used at a cut-off at 0.05. The transcript identifiers were mapped to Human Genome Organisation (HUGO) gene symbols which were used for annotation. Pathway analysis on the individual datasets was performed using TopGO package^35^, and significance threshold (adjusted p-value of < 0.01, Benjamini-Hochberg) of identified pathways was tested using Kolmogorov-Smirnov test and elim method. Identified GO terms were reduced and summarised using rrvgo package^36^.

### Discordance-Concordance Analysis

For a transcript to be considered for each *in-vitro/in-vivo* contrast, it had to be significantly differentially expressed in at least one of the two comparisons. All transcripts with adj p value below 0.05 were considered in the downstream analysis, regardless of their Log_2_FC values. A significance metric, “disco.score”, was then calculated, which accounted for both the adjusted p values and log_2_ FC values associated with each transcript. This metric adjusted the effect of low-expressing genes in the downstream analysis. The metric was calculated as follows ^14^:

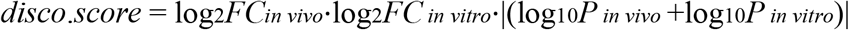

Each transcript was associated with two log_2_FC values as well as a disco.score. These were plotted on a disco plot as x and y values and colour intensity, respectively. Discoplots were divided into four quadrants which designated their concordance or discordance status. Tmod package^37^ was used to apply Gene Set Enrichment Analysis^38^ to each of the four quadrants. The resulting gene modules were summarised into parent terms and visualised on dotplots. Gene module count was visualised using heatmaps.

### *In silico* cell deconvolution

Cell fraction proportions of 22 leucocyte cell types were estimated using the online tool CibersortX^39^. The cell types under investigation were B cells naïve, B cells memory, Plasma cells, T cells CD8, T cells CD4 naïve, T cells CD4 memory resting, T cells CD4 memory activated, T cells follicular helper, T cells regulatory (Tregs), T cells gamma delta, NK cells resting, NK cells activated, Monocytes, Macrophages M0, Macrophages M1, Macrophages M2, Dendritic cells resting, Dendritic cells activated, Mast cells resting, Mast cells activated, Eosinophils and Neutrophils. The proportions were used to calculate a FC between the control and disease groups for all cell types. Cell type pairs were found between each *in vitro-in vivo* comparison and analogue to a disco.score was calculated using the *in vitro* and *in vivo* log_2_FC and their associated p-values. These metrics were plotted on cell type discoplots.

## Results

### Distinct patterns of differential transcript expression across the three data sets

Differential expression analysis for the microarray data was conducted at the transcript level using the limma package^33^. After preprocessing and visualisation using principal component analysis (Supplementary Fig 1), transcript expression data was analysed using a linear model approach^33^, for the *in vitro* infected versus uninfected control groups and the natural infection TB disease vs LTBI groups. The uninfected control and the LTBI groups served as the references. It was previously reported that there were no observable differences in whole blood transcriptome between uninfected healthy controls and those with LTBI (unless stimulated)^40^.

Firstly, for the *in vitro* dataset we conducted differential expression analysis for the 6 h, 24 h, 48 h, 72 h and 96 h *Mtb* infected timepoints against their respective uninfected control (Supplementary Fig 2). There were 45 significantly differentially expressed (SDE) genes at 6 h, 1639 at 24 h, 1409 at 48 h, 1210 at 72 h and 1623 at 96 h, with adjusted p value < 0.5 and log_2_FC threshold of 0.5. For the human natural infection datasets, we compared TB disease vs LTBI for the adult and paediatric sets separately. 2268 genes were found SDE in the paediatric comparison, and 1950 were found SDE in the adult TB comparison (Supplementary Fig 2a). Downregulation dominated the *in vitro* response at 24 h-post infection, with 65.2% (1068/1639) of all significantly differentially expressed (SDE) transcripts under-expressed (Supplementary Fig 2b). Reduced expression remained consistent across the later time points, with 63.6% (896/1409) of SDE transcripts under-expressed at 48 h (Supplementary Fig 2c) and 69.2% (837/1210) of SDE transcripts under-expressed at 72 h (Supplementary Fig 2d). In contrast, most transcripts in *in vivo* TB disease were over-expressed, with 66.0% (1499/2268) SDE transcripts upregulated in paediatric TB (Supplementary Fig 2e) and 62.6% (1221/1950) of SDE transcripts upregulated in adult TB (Supplementary Fig 2f) (exact binomial test p-value < 2.2 × 10^−16^).

Genes that were significantly upregulated and downregulated in infected patients compared to the controls were subjected to Gene Ontology (GO) pathway analysis using the topGO package^35^. Pathways enriched in downregulated genes *in vitro* (24 – 96 h) included immune activation (elim KS = 0.0052), antigen processing (elim KS = 0.0132) and defence response to bacteria (elim KS = 7.2×10^−05^), whereas pathways enriched in over-expressed genes were implicated in lymphocyte chemotaxis (elim KS = 4.2×10^−07^), neutrophil chemotaxis (elim KS = 2.3×10^−06^), and cellular response to IL-1 (elim KS = 4.0×10^−05^).

Pathways enriched in genes downregulated in natural infection included negative regulation of cytokine production (elim KS = 0.0051), cytoskeleton organisation (elim KS = 0.00606), Fc receptor signalling pathway (elim KS = 0.00580). Pathways enriched in genes upregulated in natural infection included defence response to bacteria (elim KS = 0.00022), innate immune response in mucosa (elim KS = 3.9×10^− 5^) and positive regulation of IL-1B (elim KS = 0.00091).

### Concordance and discordance analysis and accounting for the direction of gene regulation in the *in vitro-in vivo* comparisons

The disco.score was calculated for each pair of transcripts using the log_2_FC and p values as described in the Methods (***Figure 1b)***.The discordance-concordance plots were segmented into four quadrants in each *in vitro-in vivo* comparison; those which were concordantly upregulated (quadrant I), concordantly downregulated (quadrant III), discordantly regulated such that gene pairs were either upregulated *in vivo* while downregulated *in vitro* (quadrant II) or downregulated *in vivo* while upregulated *in vitro* (quadrant IV) (***Figure 2***). In this way, each *in vitro* time point was compared with both paediatric and adult patients (data for *in vivo-in vitro* 96 h comparison in Supplementary Fig 3). As the number of transcripts at 6 h post-*in vitro* infection vs control was small in comparison to the other time point comparisons, fewer transcripts were identified as either concordant or discordant. The top concordantly up-regulated genes in the adult *in vivo* vs *in vitro* comparison included *CARD17*, involved in the negative regulation of IL-1β production, *SOD2*, which has an antiapoptotic role against inflammatory cytokines^41^ and *CASP5*, which, conversely, is implicated in apoptosis. Some of the downregulated genes included *CCR3*, a chemokine receptor highly expressed in eosinophils and basophils and *DCANP1*, specifically expressed in dendritic cells. Discordantly regulated genes included *VpreB3*, which is thought to play role in B cell maturation, *SLAMF1*, involved in dendritic cell development, and *CCR7*, a gene coding for a receptor which activates B and T lymphocytes.

**Figure 1.**
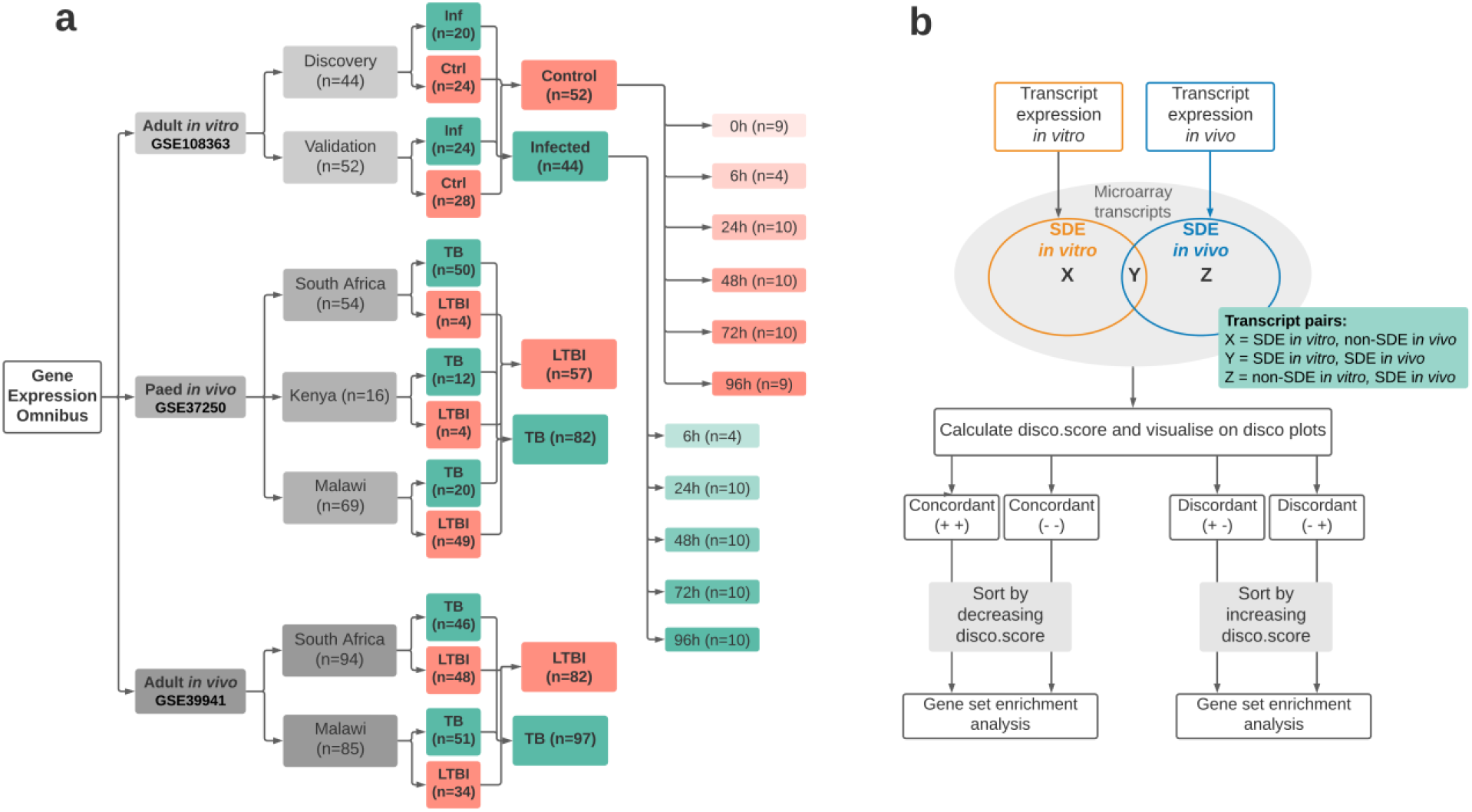
Dataset cohorts and concordance analysis design. a) Three publicly available microarray gene expression data sets deposited in the Gene Expression Omnibus were mined for the analysis. *In vitro* data derives from an adult whole blood cell infection assay; ‘Ctrl’ and ‘inf’ denote control samples (uninfected) and samples infected with *Mtb*, respectively. RNA was extracted at five time points (6 h – 96 h) post-infection in the infected and uninfected samples and at 0 h in the control/uninfected group. For *in vivo* dataset, only Human Immunodeficiency Virus (HIV)-negative and culture confirmed TB patients were included in the analysis. LTBI individuals were used as a proxy for healthy control patients. Sample size is shown for each treatment and cohort group. b) Workflow used to identify concordance and discordance between *in vitro* and *in vivo* data sets. Corresponding gene transcripts were mapped to each other to form transcript unions, composed of transcripts significant *in vivo* mapped to their non-significant counterparts *in vitro* denoted by X; transcripts significant in both data sets mapped to each other denoted by Y; transcripts significant *in vitro* mapped their non-significant counterpart *in vivo* denoted by Z. A transcript pair is said to be concordant if both of its associated log_2_FC values have the same sign. This can be further segmented into concordantly upregulated transcript pairs (both log_2_FC values are positive) and concordantly downregulated transcript pairs (both log_2_FC values are negative). A transcript pair is said to be discordant if its associated log_2_FC values have differing signs, also resulting in two final groups (positive log_2_FC *in vitro &* negative log_2_FC *in vivo;* and negative log_2_FC *in vitro &* positive log_2_FC *in vivo*). Disco.score was calculated for each corresponding transcript pair, and its value was proportional to the magnitude of concordance or discordance of said transcript pairs. Concordant transcript pairs were sorted by decreasing disco.score while discordant transcript pairs were sorted by increasing disco.score. Gene Set Enrichment Analysis (GSEA) was performed on each of the four groups of transcript pairs.

**Figure 2.**
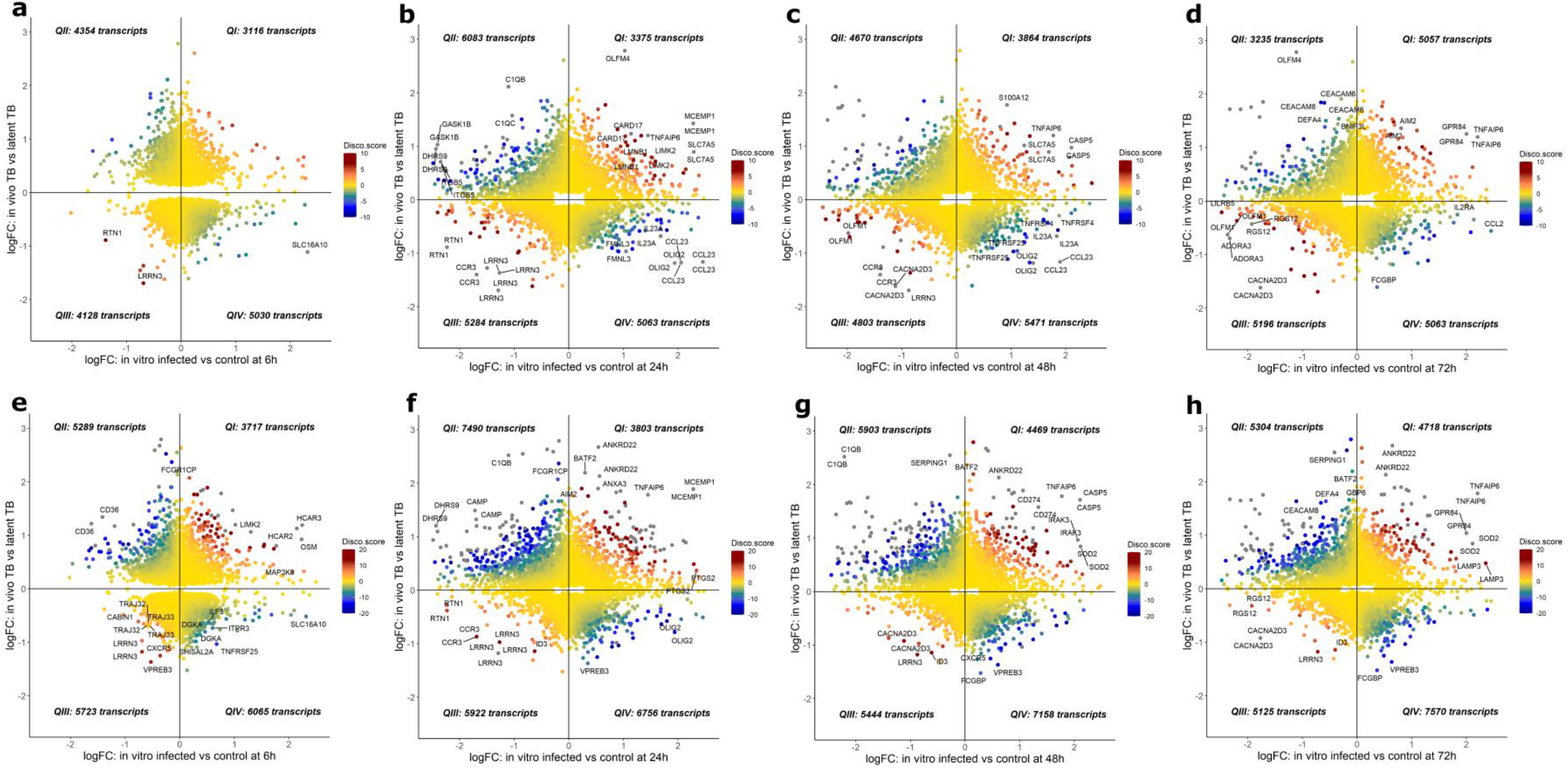
Disco plots. a-d. Evaluation of concordance of gene expression changes in adult TB and *in vitro* time series (a = 6 h, b = 24 h, c = 49 h and d = 96 h). Disco.score shown by colour intensity, ranging from blue (low concordance) to dark red (high concordance). Transcripts found in each disco plot are segmented into four quadrants (Q) as follows; QI - concordantly upregulated, QII -discordant, upregulated *in vivo &* downregulated *in vitro*, QIII -concordantly downregulated and QIV -discordant, downregulated *in vivo* and upregulated *in vitro*. Numbers of identified transcripts shown in each quadrant. e-h. Evaluation of concordance of gene expression changes in paediatric TB and *in vitro* time series (e = 6 h, f = 24 h, g = 48 h, h = 72 h).

For each *in vitro-in vivo* comparison, Gene Set Enrichment Analysis (GSEA) was performed on genes from each of the quadrants and counts of the identified gene sets (gene modules) were plotted as a heatmap, shown in ***Figure 3***. Heatmaps revealed that there is a trend towards concordance in earlier time points (0 – 48 h), while discordance is more dominant from 48 h onwards. Paediatric *in vivo-in vitro* showed more concordantly downregulated modules (***Figure 3c***), while adult *in vivo-in vitro* has a distinctly different pattern which showed many more concordantly upregulated modules (***Figure 3a***). The largest number of modules were identified as discordantly regulated, upregulated *in vitro* and downregulated *in vivo*.

**Figure 3.**
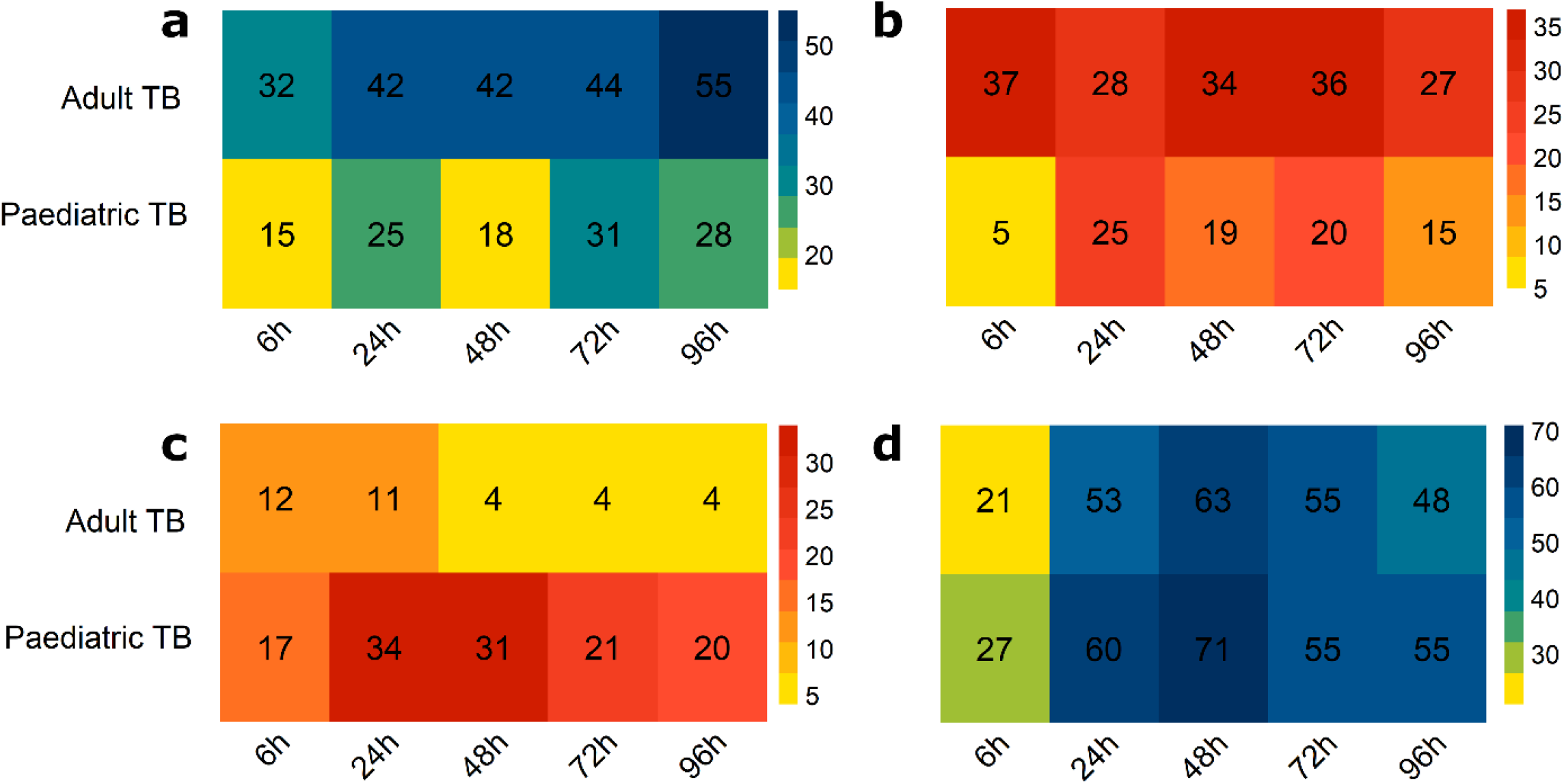
Counts of concordantly and discordantly regulated gene modules. Gene Set Enrichment Analysis (GSEA) was performed on transcripts from the four disco plot quadrants for each *in vitro-in vivo* comparison. Identified modules were counted and plotted as heatmaps in the same arrangement as quadrants of a discoplot from which the transcripts were taken. Colour intensity ranging from yellow to red for concordant modules, and yellow to blue for discordant modules, denotes increasing number of modules identified. a) Counts of discordant modules, upregulated *in vivo*, and downregulated *in vitro*. b) Counts of concordantly upregulated modules. c) Counts of concordantly downregulated modules. d) Counts of discordant modules, downregulated *in vivo*, and upregulated *in vitro*.

### Concordance and discordance analysis of the adult *in vivo-in vitro* comparison

To explore the identified gene modules in a more comprehensive way, gene modules describing similar biological pathways were summarised, and thereafter visualised using dot plots shown in ***Figure 4*** and ***Figure 5***. Remarkably, time signatures of concordantly and discordantly regulated modules can be clearly distinguished in these plots.

**Figure 4.**
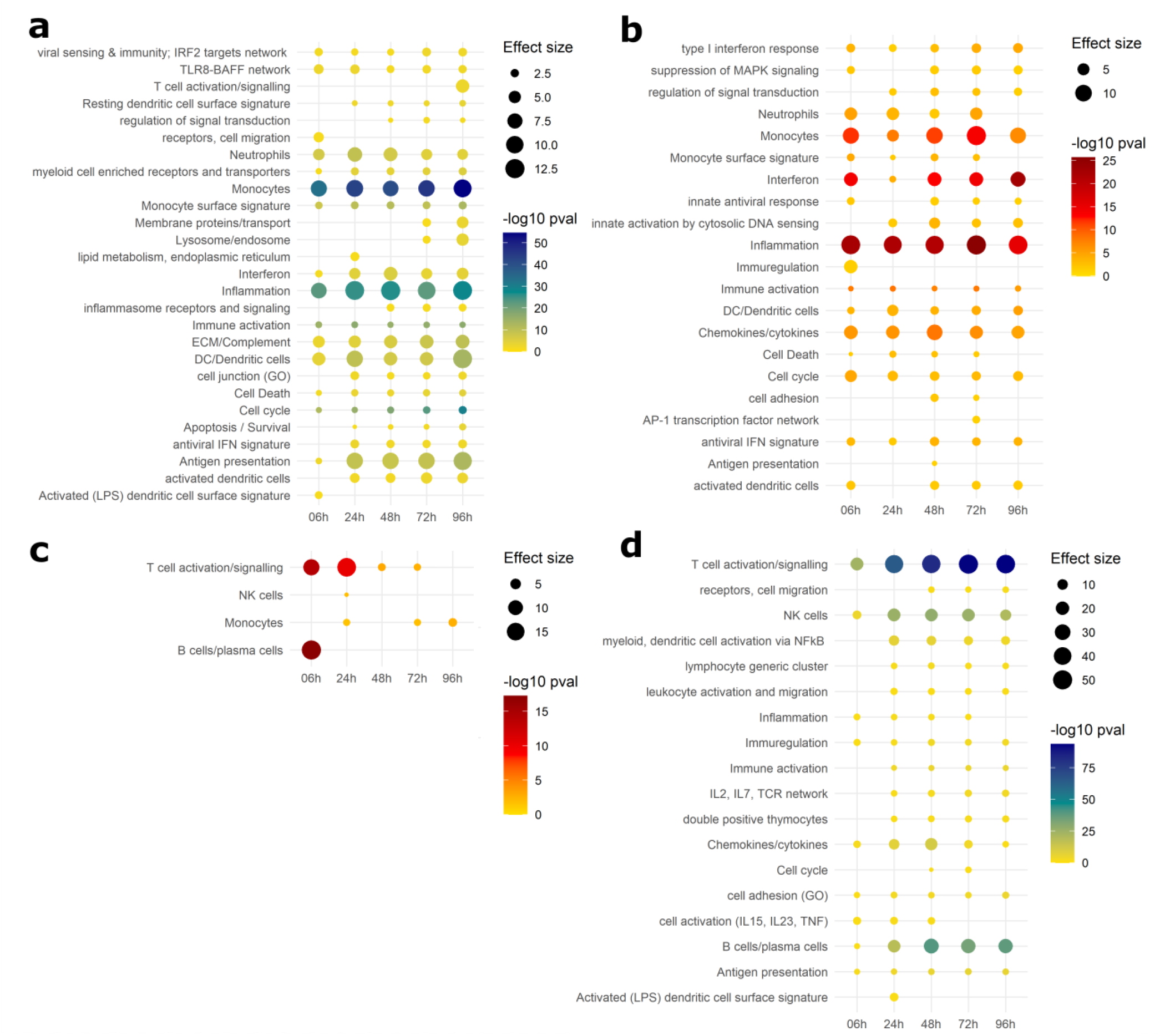
Dotplot of gene modules identified in adult *in vivo-in vitro* comparison. Gene Set Enrichment Analysis (GSEA) was performed on transcripts from the disco plot quadrants. Similar modules were summarised and plotted such that p-value of module enrichment is illustrated by the intensity of the colour (yellow – blue gradient denotes increasing significance of discordant pathways, yellow-red gradient denotes increasing significance of concordant pathways) and the effect size by the size of the dot. a) Gene modules upregulated *in vivo* and downregulated *in vitro*. b) Concordantly upregulated gene modules. c) Concordantly downregulated gene modules. d) Gene modules downregulated *in vivo* and upregulated *in vitro*.

**Figure 5.**
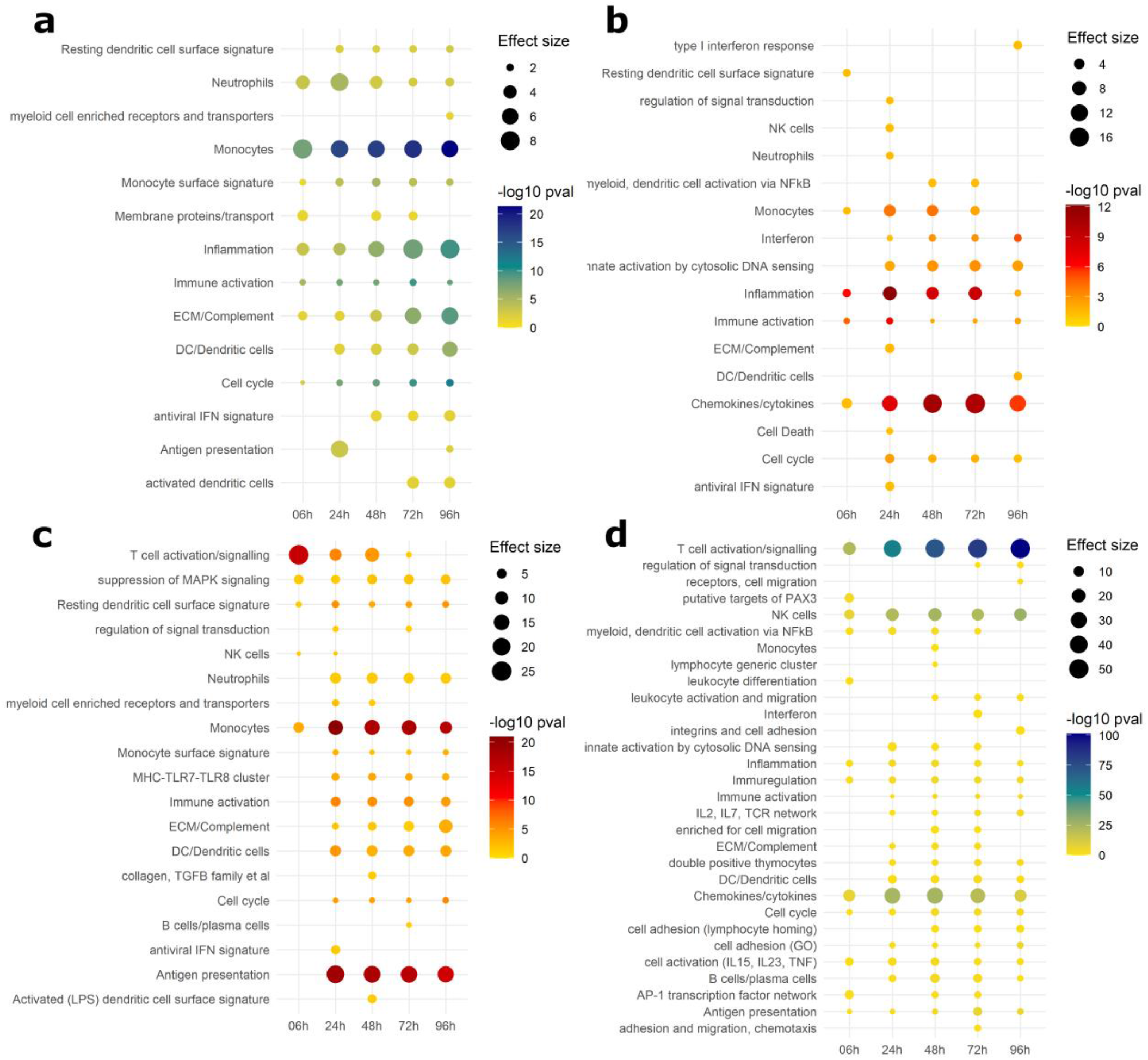
Dotplot of gene modules identified in paediatric *in vivo-in vitro* comparison. Gene Set Enrichment Analysis (GSEA) was performed on transcripts from the disco plot quadrants. Similar modules were summarised and plotted such that p-value of module enrichment is illustrated by the intensity of the colour and the effect size by the size of the dot. a) Gene modules upregulated *in vivo* and downregulated *in vitro*. b) Concordantly upregulated gene modules. c) Concordantly downregulated gene modules. d) Gene modules downregulated *in vivo* and upregulated *in vitro*.

In agreement with our previous findings, concordance of the adult *in vivo*-*in vitro* comparison was primarily seen in upregulated modules (***Figure 4b***, representing quadrant I), whereas very few concordantly downregulated modules were identified (***Figure 4c***, representing quadrant III). Concordant upregulation (***Figure 4b***) was driven by inflammation exhibiting the highest effect size (average cES = 12.8, average p val = 4.36×10^−22^) across all timepoints, as well as pathways relating to monocyte enrichment (average cES = 9.34, average p val = 4.57×10^−11^), chemokines (average cES = 6.67, average p val = 4.7×10^−7^), and cytokines (type 1 interferon response) (average cES = 2.98, average p val = 4.6×10^−4^). T cell and B cell modules were strongly concordantly downregulated in the comparisons with 6 – 24 h *in vitro* (cES = 14.5, p val = 6.1×10^−13^ and cES = 17.95, p val = 5.09×10^−18^ respectively) after which the abundance and significance of these pathways rapidly decreased. Concordant downregulation of monocyte-related pathways was also discernible, however the number of genes the modules were enriched in is 7-fold lower than T cell modules.

***Figure 4d*** (representing quadrant IV) summarises modules that are upregulated *in vitro* and downregulated *in vivo*. Here, the discordance is most apparent in T cell and B cell modules, where it increases over time (T cells at 6 h: cES = 17.54, p val = 8.27×10^−25^, at 96 h: cES = 46.87, p val = 1.67×10^−94^. B cells at 6 h: cES = 2.13, p val = 0.025, at 96 h: cES = 23.5, p val = 8.56×10^−39^). NK cells also have a slight time signature, with the strongest signal at 24-72 h (cES = 17, p val = 2.3×10^−27^). Other modules regulated in this way also relate to lymphocytes, their activation and function.

In ***Figure 4a*** (quadrant II), monocyte and inflammation modules can also be found (average cES = 9.09, p val = 2.87×10^−44^ and average cES = 11.08, p val = 6.38×10^−26^ respectively), alongside dendritic cell and antigen presentation modules.

### Concordance analysis of the paediatric *in vivo-in vitro* comparison

GSEA was done on genes from the individual quadrants of disco plot comparisons of paediatric *in vivo-in vitro* time series, as before. ***Figure 5b*** shows concordantly upregulated modules (quadrant I), which are dominated by a chemokine/cytokine module (average cES = 11.2, average p value = 7.43×10^−8^), and secondary to it, inflammation, and monocyte related modules (cES = 5.09, p val = 3.01×10^−8^ and cES = 4.07, 1.28×10^−3^ respectively). Many more concordantly downregulated modules (quadrant III) were identified in ***Figure 5c***, compared to the adult *in vivo-in vitro* comparison. These include T cell activation, greatest at 6 h (cES = 25.4, p val = 3.1×10^−16^) and decreasing in both effect size and significance at 48 h (cES = 11.5, p val = 1.62×10^−5^). Modules relating to monocytes and antigen presentation also drove the concordance of downregulation in this comparison.

As in the adult *in vivo-in vitro* comparison, monocytes were also identified in quadrant II, i.e., modules that were upregulated *in vivo* and downregulated *in vitro* (***Figure 5a***). Along with inflammation and dendritic cells, these modules have increasing signal towards later time points.

Modules that are upregulated *in vitro* and downregulated *in vivo* in ***Figure 5d*** (Quadrant IV) are primarily related to T cell signalling as in adult *in vivo-in vitro* comparison; however, B cell modules were identified but less pronounced (cES = 5.24, p val = 0.003). Other modules here included NK cells and those related to regulating chemokines and cytokines.

### Comparison of *in silico* cell deconvolution corroborates the findings of the concordance analysis

Transcript expression data was also used for *in silico* cell deconvolution (CibersortX^39^) and the output of relative cell fractions of 22 immune cell types was split into adaptive and innate cells. The differences between TB disease and LTBI controls in adult and paediatric patients (*in vivo* data) are largely consistent across innate cells (***Figure 7b*** and ***c***) and adaptive cells (***Figure 6b*** and ***c)***. The only exceptions are in T cells - CD4 memory activated, - CD4 memory resting and T cells regulatory fractions which are significantly elevated, reduced and reduced, respectively, in the adult TB and control comparison, but do not show any significant differences in the paediatric TB versus LTBI groups.

**Figure 6.**
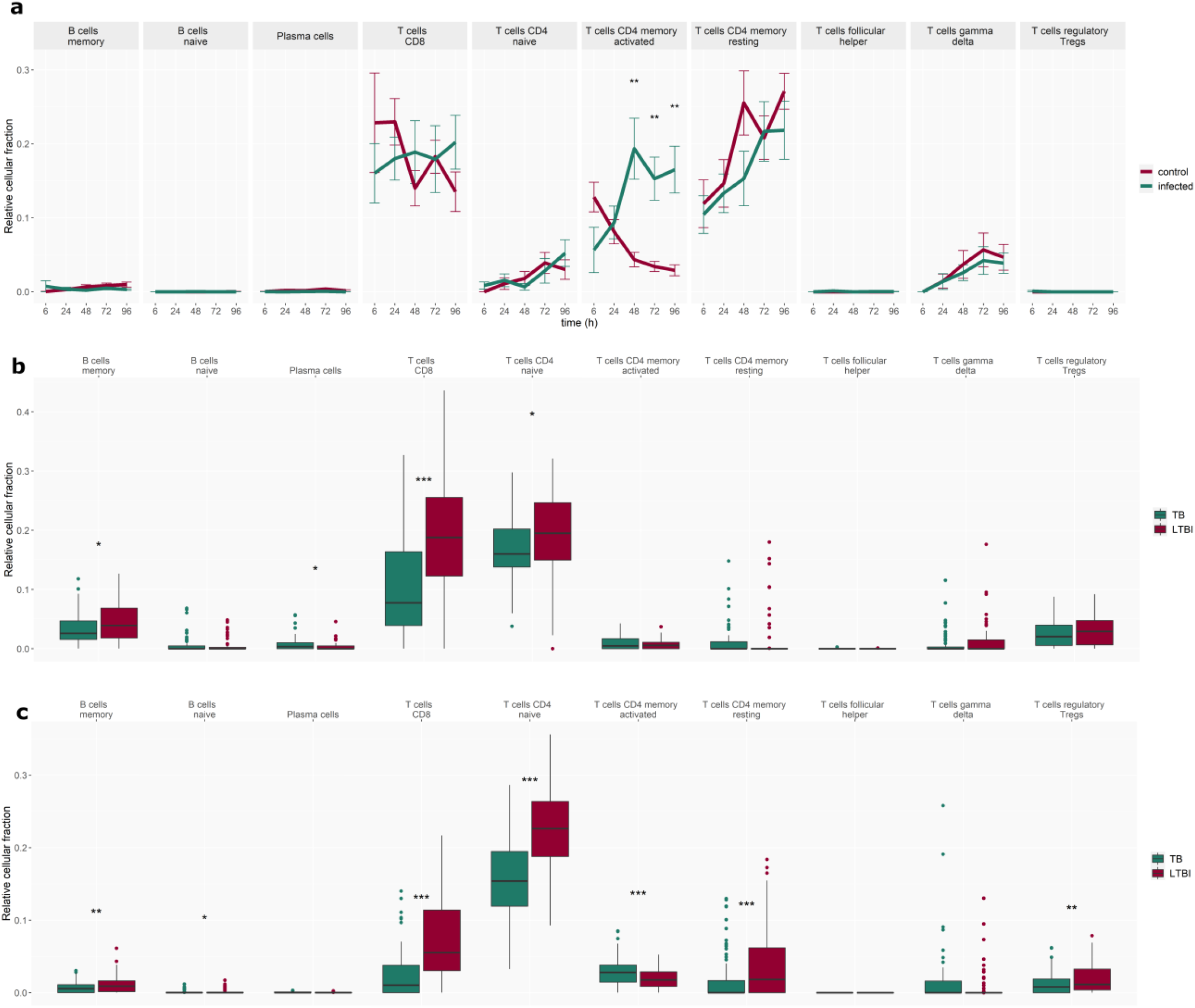
Cell deconvolution using CibersortX – adaptive cells. Changes in adaptive cell fractions a) *in vitro* over time in the *Mtb*-infected and control group and *in vivo* in b) paediatric TB patients and c) adult TB patients and corresponding LTBI control patient groups. Error bars depict standard deviation from the mean, *p<0.05, **p<0.005, ***p<0.001.

**Figure 7.**
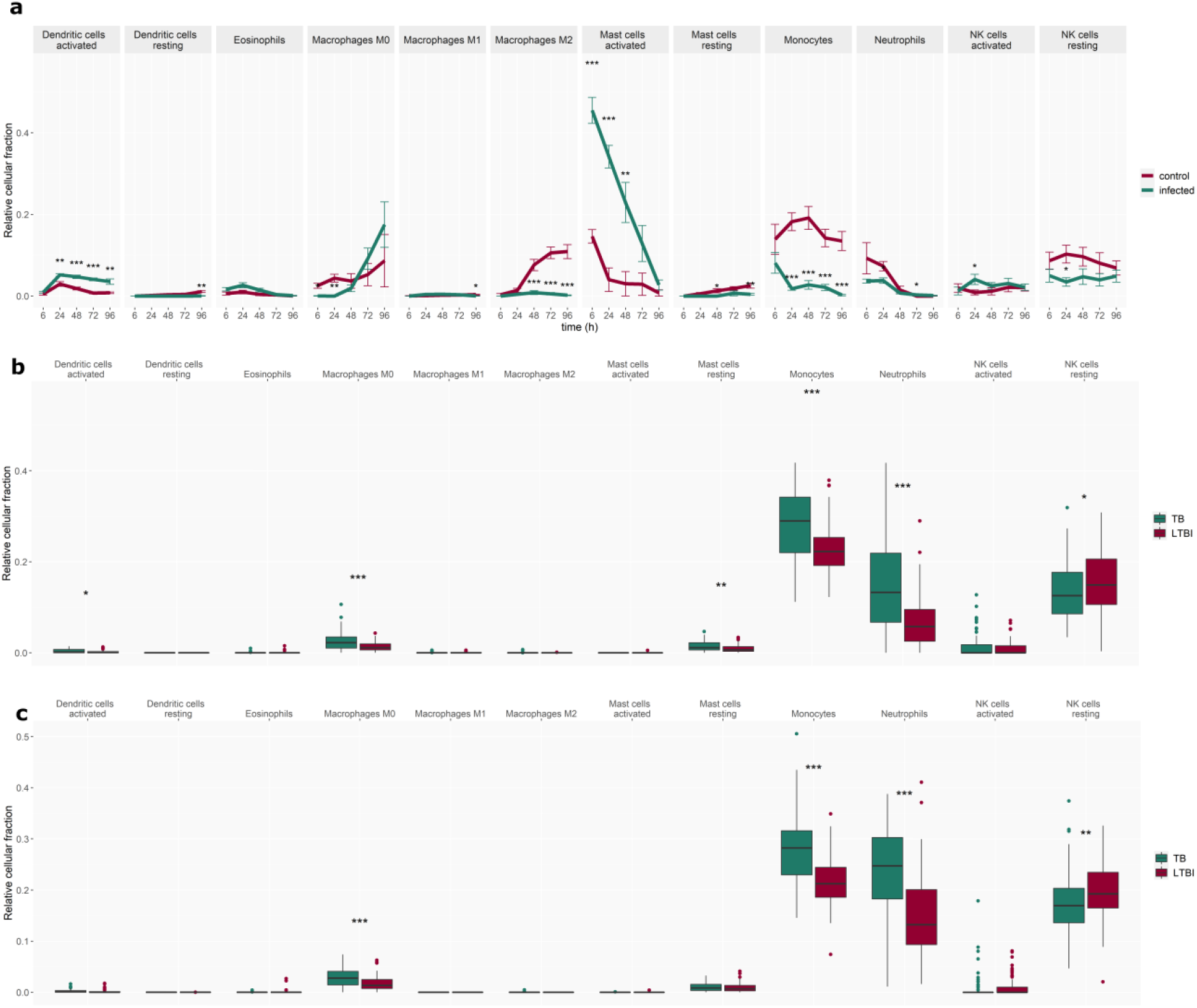
Cell deconvolution using CibersortX – innate cells. Changes in innate cell fractions a) *in vitro* over time in the *Mtb*-infected and control group and *in vivo* in b) paediatric TB patients and c) adult TB patients and corresponding LTBI control patient groups. Error bars depict standard deviation from the mean, *p<0.05, **p<0.005, ***p<0.001.

No significant changes over time were identified in the cell fractions *in vitro* between control and TB-infected with the exception of CD4 memory activated fraction, which increases from 24 h onwards post-infection.

Indeed, in ***Figure 7*** monocyte, and neutrophil fractions are elevated *in vivo*, while they are diminished *in vitro* from the early time points onward. Mast cells account for the biggest fraction *in vitro* at 6 h post-infection, but rapidly decrease to the level of the control at later time points. *In vitro-in vivo* agree in dendritic cell fractions as well as non-activated (M0) macrophages which are elevated from 24 h and 48 h, respectively.

## Discussion

Reproducible and clinically translatable TB research is crucial to address the urgent need for vaccines, novel therapeutics, and more sensitive diagnostic tests. *In vitro* studies combining WBA with gene expression readouts have elucidated important facets of the interplay between *Mtb* and the host, particularly regarding immune cell interactions ^6,16,42^. WBA contain most immune cell types and thus account for a much broader and a more physiological representation of the immune system than other *in vitro* cell type specific infection models. Interpreting and extrapolating *in vitro* observations from the WBA requires knowledge of the similarities and differences between the WBA and specific elements of natural TB infection and disease. For example, differences in cell populations resulting from longer culture and infection treatment times may drive divergence between *in vitro* and *in vivo* systems, if more short-lived cell types, such as neutrophils^43^, have undergone apoptosis and cannot participate in the full response to the pathogen. No studies have been performed to elucidate how representative the *Mtb* infection WBA transcriptomic profiles are in comparison to natural human infection and disease.

In this study we characterised the concordance and discordance of gene expression profiles between an *in vitro Mtb* infection assay which uses peripheral blood from healthy adult donors and *in vivo* peripheral circulating blood samples from paediatric and adult TB patients. We compared the transcriptomic profiles and cell fractions between WBA and the patient data and identified conserved and divergent host immune gene expression. To better understand how the WBA relates to *in vivo* TB disease, and particularly how different experimental time points of the assay relate to specific elements of the host immune response, we implemented a concordance analysis of gene regulation accompanied by GSEA and *in silico* cell deconvolution^39^. The *in vitro* study compared TB infection vs uninfected controls, whereas the *in vivo* studies compared TB disease vs LTBI. Thus, the patient datasets have captured systemic disease aspects: lung pathology, tissue necrosis, unwellness in addition to the *Mtb* immune escape mechanisms which enabled TB disease to develop. This study has enabled us to assess which elements of the *in vitro* model recapitulate aspects of the *in vivo* anti-mycobacterial immune response.

We observed that concordance between *in vivo-in vitro* data sets is primarily associated with immune activation for adult TB disease and immune suppression for paediatric TB disease. Concordance in the adult *in vivo* versus WBA comparison was driven by upregulated modules including those involved in inflammation, pro-inflammatory signalling, neutrophil, and monocyte enrichment, which are important for the recognition of *Mtb* infection and the initiation of a large-scale immune response to the pathogen^7^. Fewer concordantly downregulated modules were observed, including those involved in T and B cell function, with the concordance identified at earlier time points (0 h and 24 h). We also identified the concordant and discordant gene modules in the WBA dataset that were consistent in both adult and paediatric *in vivo* TB disease datasets. We found that both comparisons had common concordance in modules related to inflammatory pathways expected during an infection with mycobacteria, such as cytokine (particularly IFN signatures) and chemokine signalling, and modules relating to T cell suppression and monocytes, most notably in earlier time points (6 – 24 h). Common discordant modules dominated both comparisons with the later time points of the WBA (72 – 96 h), especially those relating T cell signalling, dendritic cell activation and monocytes.

Apoptotic depletion of macrophages and dendritic cells in the *in vitro* model without the means to replenish them will lead to a substantial divergence between the WBA and *in vivo* studies. Therefore, the higher abundance of T cell signatures at later time points (48 – 96 h) *in vitro* relative to *in vivo* may be a result of a higher fraction in the bulk gene expression analysis following the loss of other cell types. Delay of the onset of the adaptive immune response *in vivo* as well as T cell exhaustion, manifested by reduced T-cell activation, may also explain the large contrast between the *in vivo* and *in vitro* systems. As mentioned previously, we also observed concordance in downregulated T cell modules in both paediatric and adult *in vivo-in vitro* comparisons at 6 – 24 h. It should be noted that, while the *in silico* cell deconvolution analysis has been shown to accurately predict the proportion of CD4 and CD8 T cell fractions in previous studies, estimations of other T cell fractions may be less accurate based on previous validation studies using this package^44^ and a more specific analysis of T cell fractions (e.g. Th1, Th2, and Th17 cells) may provide further insights towards understanding the concordance and discordance results observed here. Furthermore, the differences between paediatric and adult T cell populations and T cell priming are an important consideration when analysing these data. However, the prediction of the certain T cell populations which are known to differ between these groups (for example γδ T cells are highly relevant for paediatric TB responses^45^) may not be possible with many of the current *in silico* deconvolution tools.

The abundance of discordant modules increased at later time points, driven particularly by gene modules related to inflammation, T cells, monocytes, and dendritic cells. In both adult and paediatric *in vivo* vs *in vitro* comparisons, discordant modules based on genes up-regulated *in vivo* and down-regulated *in vitro* were associated with enrichment in monocytes, inflammation, and antigen presentation. On the other hand, modules based on genes down-regulated *in vivo* and up-regulated *in vitro* are primarily involved in T cell and NK cell regulation. These findings are largely supported by *in silico* cell deconvolution which revealed that the monocyte cell fraction significantly decreases in the infected group after 24 h *in vitro*, while T cell fractions stay constant (CD8) throughout the assay timeline or indeed increase in abundance (CD4) towards the later time points. In contrast, in the *in vivo* cell deconvolution results, the monocytes fraction is significantly elevated in both adult and paediatric TB patients. M0 macrophages, which derive from monocytes, were also found to be significantly elevated in both TB patient datasets. In contrast, CD8 and CD4 naïve T cells are significantly decreased in TB patients from both datasets. However, in the adult patient cohort, activated CD4 memory T cells are increased, while resting CD4 memory T cells were decreased.

When comparing the WBA versus the paediatric *in vivo* dataset, we identified gene modules involved in host immune signalling, such as those related to chemokines, cytokines and other markers of inflammation, as concordantly upregulated, while modules associated with monocytes and dendritic cells, which harbour *Mtb* proliferation, were concordantly downregulated along with antigen presentation and immune activation signatures. This observed downregulation may reflect *Mtb* suppression of the host immune response, which has been previously described in paediatric TB disease^46,47^. By contrast, in the adult comparison antigen presentation was discordantly upregulated in the adult *in vivo* patient dataset and downregulated in the WBA dataset. Differences in the whole blood transcriptome in children and adults with TB disease have been previously reported (e.g., inhibition of neutrophil degranulation and difficulty in T cell priming^47,48^). It is worth noting that the WBA utilises whole blood from naïve adult donors, and age difference between the WBA and the *in vivo* paediatric dataset could be driving some of the differences reported. In addition, the sample size for the groups in the paediatric *in vivo* dataset is lower compared to that in the adult *in vivo* dataset, which may be the cause of the slightly lower p-value significance levels of the differentially expressed transcripts in the paediatric cohort.

The study has certain limitations. As the WBA uses blood from healthy adult donors, some of the differences observed in the comparisons encompassing the paediatric dataset may be attributed to age. Conducting a WBA-based study that uses blood from children may be more relevant in elucidating paediatric-specific host response elements^49^. Secondly, although previous studies have reported the lack of differences between the transcriptomic profiles in unstimulated blood of Interferon-Gamma Release Assay (IGRA) positive individuals and healthy controls, discordance reported between the *in vitro* and *in vivo* datasets could be attributed to differences in the baseline comparator group (LTBI *in vivo* vs uninfected blood *in vitro)*. The analysis needs to be coupled with other readouts, such as experimental flow cytometry, in order to disentangle the causality of the relationships between the gene modules. Lastly, our analysis was conducted on bulk microarray gene expression data, which profiles transcripts included on the array, rather than the whole transcriptome. Future studies can be expanded to RNA-sequencing data and especially single-cell RNA-sequencing, to further examine the complexity of host immune response to *Mtb*, including multiple single cellular populations within a system.

In summary, we present an adaptation of concordance analysis for the comparison of host immune response *in vivo* and *in vitro*. Our findings suggest that earlier time points (24 – 48 h) of the whole blood assay are more concordant with patient disease, reflected by both the gene expression and *in silico* cell deconvolution at these time points. There are specific similarities of the assay with paediatric and adult *in vivo* disease, such as upregulation of inflammatory signatures, particularly relating to IFN signalling, providing a reference to tailor the assay to paediatric and adult TB disease in future studies. Our study also provides valuable information about immune cell components driving discordance between the datasets, which manifest especially at later time points (72 – 96 h). Understanding the limitations of *in vitro* assays is vital for its continued use and consequently for conducting more robust and translatable research. Identifying the magnitude and source of concordance and discordance of bulk RNA expression profiling in *in vitro* WBA compared to *in vivo* natural infection can help better understand and tailor the *in vitro* models to explore relevant biological questions of interest.

## Data availability

Raw data is available at Gene Expression Omnibus database https://www.ncbi.nlm.nih.gov/geo/. Accession numbers for the datasets are as follows; *in vitro* dataset: GSE108363, adult *in vivo* dataset: GSE39941, paediatric *in vivo* dataset: GSE37250.

Scripts are available on request.

## Competing interests

Authors declare no competing interests.

## Acknowledgements

We would like to thank the authors of the manuscripts for making the datasets used in this study publicly available. C.B. acknowledges support from the NIHR Imperial College BRC (Imperial 4i fellowship-RDA02). M.K. acknowledges support from The Wellcome Trust and the Medical Research Foundation grants (206508/Z/17/Z and MRF-160-0008-ELP-KAFO-C0801). P.B., A.C., C.B., S.M.N, M.L. and M.K. acknowledge support from the NIHR Imperial College BRC.

## Author contributions

A.C. and M.K.: Conceptualization, P.B., A.C. and M.K.: Methodology, P.B.: Data curation, P.B., A.C. and M.K.: Writing-original draft preparation. P.B., A.C. and M.K: Visualization, A. C., S.M.N., M.L., M.K, Supervision, All authors: Writing, reviewing and, editing

## Notes

### Competing Interest Statement

The authors have declared no competing interest.

### Summary of Updates

to correct for pdf conversion formatting errors

## References

1. GLOBAL TUBERCULOSIS REPORT 2021. (2021).

2. Cruz-Knight, W. & Blake-Gumbs, L. Tuberculosis: An Overview. Primary Care: Clinics in Office Practice 40, 743–756 (2013).

3. Houben, R. M. G. J. & Dodd, P. J. The Global Burden of Latent Tuberculosis Infection: A Reestimation Using Mathematical Modelling. PLoS Medicine 13, (2016).

4. Flynn, J. L. & Chan, J. Tuberculosis: Latency and Reactivation. Infection and Immunity 69, 4195–4201 (2001).

5. Fogel, N. Tuberculosis: A disease without boundaries. Tuberculosis 95, 527–531 (2015).

6. von Both, U. et al. Mycobacterium tuberculosis Exploits a Molecular Off Switch of the Immune System for Intracellular Survival. Sci Rep (2018).

7. de Martino, M., Lodi, L., Galli, L. & Chiappini, E. Immune Response to Mycobacterium tuberculosis: A Narrative Review. Frontiers in Pediatrics 7, 350 (2019).

8. Gliddon, H. D. et al. Identification of Reduced Host Transcriptomic Signatures for Tuberculosis Disease and Digital PCR-Based Validation and Quantification. Frontiers in Immunology 12, (2021).

9. Gliddon, H. D., Herberg, J. A., Levin, M. & Kaforou, M. Genome-wide host RNA signatures of infectious diseases: discovery and clinical translation. Immunology 153, 171–178 (2017).

10. Anderson, S. T. et al. Diagnosis of Childhood Tuberculosis and Host RNA Expression in Africa. New England Journal of Medicine 370, 1712–1723 (2014).

11. Hoang, L. T. et al. Transcriptomic signatures for diagnosing tuberculosis in clinical practice: a prospective, multicentre cohort study. The Lancet Infectious Diseases 21, 366–375 (2021).

12. Mulenga, H. et al. Performance of host blood transcriptomic signatures for diagnosing and predicting progression to tuberculosis disease in HIV-negative adults and adolescents: a systematic review protocol. BMJ Open 9, (2019).

13. Sweeney, T. E., Braviak, L., Tato, C. M. & Khatri, P. Genome-wide expression for diagnosis of pulmonary tuberculosis: a multicohort analysis. The Lancet Respiratory Medicine 4, 213–224 (2016).

14. Domaszewska, T. et al. Concordant and discordant gene expression patterns in mouse strains identify best-fit animal model for human tuberculosis. Scientific Reports 7, (2017).

15. Whatney, W. E. et al. A High Throughput Whole Blood Assay for Analysis of Multiple Antigen-Specific T Cell Responses in Human Mycobacterium tuberculosis Infection. The Journal of Immunology 200, 3008–3019 (2018).

16. Newton, S., Martineau, A. & Kampmann, B. A Functional Whole Blood Assay to Measure Viability of Mycobacteria, using Reporter-Gene Tagged BCG or M.Tb (BCG lux/M.Tb lux). Journal of Visualized Experiments (2011) doi:10.3791/3332.

17. Silva, D., Ponte, C. G. G., Hacker, M. A. & Antas, P. R. Z. A whole blood assay as a simple, broad assessment of cytokines and chemokines to evaluate human immune responses to Mycobacterium tuberculosis antigens. Acta Tropica 127, 75–81 (2013).

18. R Core Team. R: A Language and Environment for Statistical Computing. (2020).

19. Kaforou, M. et al. Detection of Tuberculosis in HIV-Infected and -Uninfected African Adults Using Whole Blood RNA Expression Signatures: A Case-Control Study. PLoS Medicine 10, (2013).

20. Leek, J. T. et al. sva: Surrogate Variable Analysis. (2020).

21. Blighe, K. & Lun, A. PCAtools: PCAtools: Everything Principal Components Analysis. (2020).

22. Kolde, R. pheatmap: Pretty Heatmaps. (2019).

23. Wickham, H. ggplot2: Elegant Graphics for Data Analysis. (Springer-Verlag New York, 2016).

24. Slowikowski, K. ggrepel: Automatically Position Non-Overlapping Text Labels with “ggplot2.” (2021).

25. Bache, S. M. & Wickham, H. magrittr: A Forward-Pipe Operator for R. (2020).

26. Neuwirth, E. RColorBrewer: ColorBrewer Palettes. (2014).

27. Ewing, M. mgsub: Safe, Multiple, Simultaneous String Substitution. (2020).

28. Dowle, M. & Srinivasan, A. data.table: Extension of ‘data.frame’. (2021).

29. Durinck, S. et al. BioMart and Bioconductor: a powerful link between biological databases and microarray data analysis. Bioinformatics 21, 3439–3440 (2005).

30. Friedman, A. B. taRifx: Collection of Utility and Convenience Functions. (2020).

31. Wickham, H. Reshaping data with the reshape package. Journal of Statistical Software 21, (2007).

32. Robinson MD, McCarthy DJ & Smyth GK. edgeR: a Bioconductor package for differential expression analysis of digital gene expression data. Bioinformatics 26, 139–140 (2010).

33. Ritchie, M. E. et al. limma powers differential expression analyses for RNA-sequencing and microarray studies. Nucleic Acids Research 43, e47 (2015).

34. Blighe, K., Rana, S. & Lewis, M. EnhancedVolcano: Publication-ready volcano plots with enhanced colouring and labeling. (2020).

35. Alexa, A. & Rahnenfuhrer, J. topGO: Enrichment Analysis for Gene Ontology. (2020).

36. Sayols, S. rrvgo: a Bioconductor package to reduce and visualize Gene Ontology terms. (2020).

37. Weiner, J. tmod: Feature Set Enrichment Analysis for Metabolomics and Transcriptomics. (2020).

38. Subramanian, A. et al. Gene set enrichment analysis: A knowledge-based approach for interpreting genome-wide expression profiles. Proceedings of the National Academy of Sciences 102, 15545–15550 (2005).

39. Newman, A. M. et al. Determining cell type abundance and expression from bulk tissues with digital cytometry. Nature Biotechnology 37, (2019).

40. Broderick, C., Cliff, J. M., Lee, J. S., Kaforou, M. & Moore, D. A. Host transcriptional response to TB preventive therapy differentiates two sub-groups of IGRA-positive individuals. Tuberculosis 127, (2021).

41. Montano, M. A. E. et al. Inflammatory cytokines in vitro production are associated with Ala16Val superoxide dismutase gene polymorphism of peripheral blood mononuclear cells. Cytokine 60, 30–33 (2012).

42. Kampmann, B. et al. Novel Human In Vitro System for Evaluating Antimycobacterial Vaccines. Infection and Immunity 72, 6401–6407 (2004).

43. Tak, T., Tesselaar, K., Pillay, J., Borghans, J. A. M. & Koenderman, L. What’s your age again? Determination of human neutrophil half-lives revisited. Journal of Leukocyte Biology 94, 595–601 (2013).

44. Miao, Y. et al. ImmuCellAI: A Unique Method for Comprehensive T-Cell Subsets Abundance Prediction and its Application in Cancer Immunotherapy. Advanced Science 7, 1902880 (2020).

45. Whittaker, E., Nicol, M., Zar, H. J. & Kampmann, B. Regulatory T Cells and Pro-inflammatory Responses Predominate in Children with Tuberculosis. Frontiers in Immunology 8, (2017).

46. Hemingway, C. et al. Childhood tuberculosis is associated with decreased abundance of T cell gene transcripts and impaired T cell function. PLoS ONE 12, (2017).

47. Bah, S. Y., Forster, T., Dickinson, P., Kampmann, B. & Ghazal, P. Meta-analysis identification of highly robust and differential immune-metabolic signatures of systemic host response to acute and latent tuberculosis in children and adults. Frontiers in Genetics 9, (2018).

48. Basu Roy, R., Whittaker, E. & Kampmann, B. Current understanding of the immune response to tuberculosis in children. Current Opinion in Infectious Diseases 25, 250–257 (2012).

49. Roy, R. B. et al. An auto-luminescent fluorescent BCG whole blood assay to enable evaluation of paediatric mycobacterial responses using minimal blood volumes. Frontiers in Pediatrics 7, (2019).

