## Supplementary for "Comparative transcriptomic analysis of whole blood mycobacterial growth assays and tuberculosis patients’ blood RNA profiles"

Table of Contents

**Supplementary Table 1:** Numbers of significantly differentially expressed transcripts in individual datasets	**2**

**Supplementary Table 2:** Numbers of transcripts after mapping into transcript unions	**2**

**Supplementary Figure 1:** Dimensionality reduction of the gene expression data using principal component analysis**.**	**3**

**Supplementary Figure 2:** Differential transcript expression	**4**

**Supplementary Figure 3:** Disco plots for 96 h concordance comparisons	**5**

**Supplementary Files:** Gene Ontology pathway lists – xlsx file list	**5**

| Dataset | Significantly expressed transcripts (adj p-val < 0.05) |
| --- | --- |
| *In vitro* 6 h vs 0h | 45 |
| *In vitro* 24 h vs 0h | 2152 |
| *In vitro* 48 h vs 0h | 1767 |
| *In vitro* 72 h vs 0h | 1476 |
| *In vitro* 96 h vs 0h | 2093 |
| Paediatric *in vivo TB vs LTBI* | 13,051 |
| Adult *in vivo TB vs LTBI* | 10,007 |

**Supplementary Table 1 Numbers of significantly differentially expressed transcripts in individual datasets.**

|  | Paediatric | Adult |
| --- | --- | --- |
| *In vitro* 6 h | 16,628 | 20,794 |
| *In vitro* 24 h | 19,805 | 23,971 |
| *In vitro* 48 h | 18,808 | 22,974 |
| *In vitro* 72 h | 18,551 | 22,717 |
| *In vitro* 96 h | 19,410 | 23,576 |

**Supplementary Table 2 Numbers of transcripts after mapping into transcript unions.** Significantly expressed transcripts from *in vitro* and *in vivo* datasets were mapped to each other and used for discordance, concordance analysis.

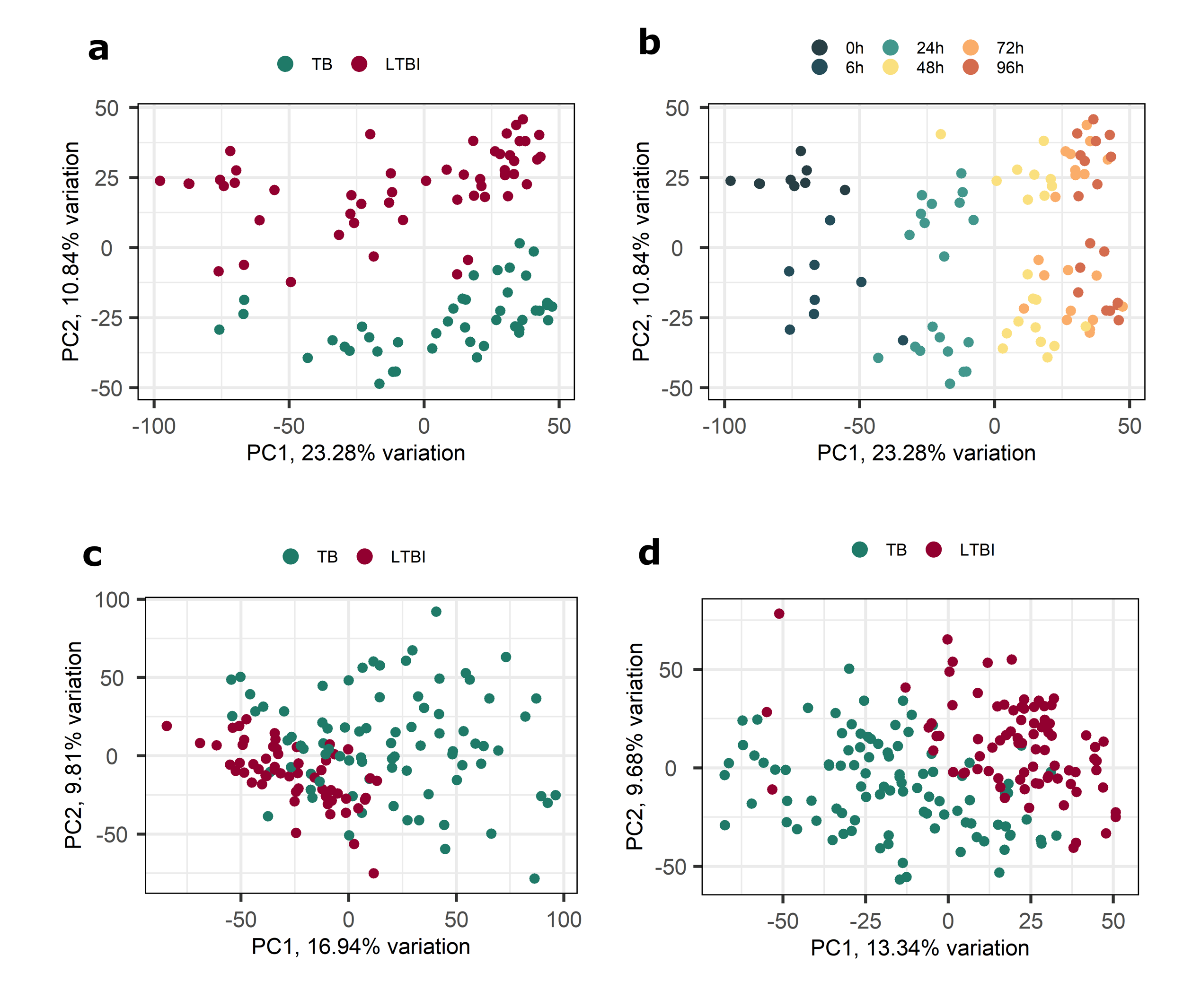

**Supplementary Figure 1: Dimensionality reduction of the gene expression data using principal component analysis.**

a) and b) show principal component analysis of *in vitro* data, coloured by infected versus respective uninfected control (
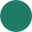
 - TB and
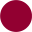
 - LTBI) at different time points (
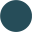
 - 6 h,
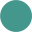
 - 24 h,
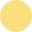
 - 48 h,
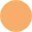
 - 72 h,
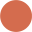
 - 96 h), respectively. c) and d) show principal component analysis of paediatric and adult *in vivo* data, respectively.

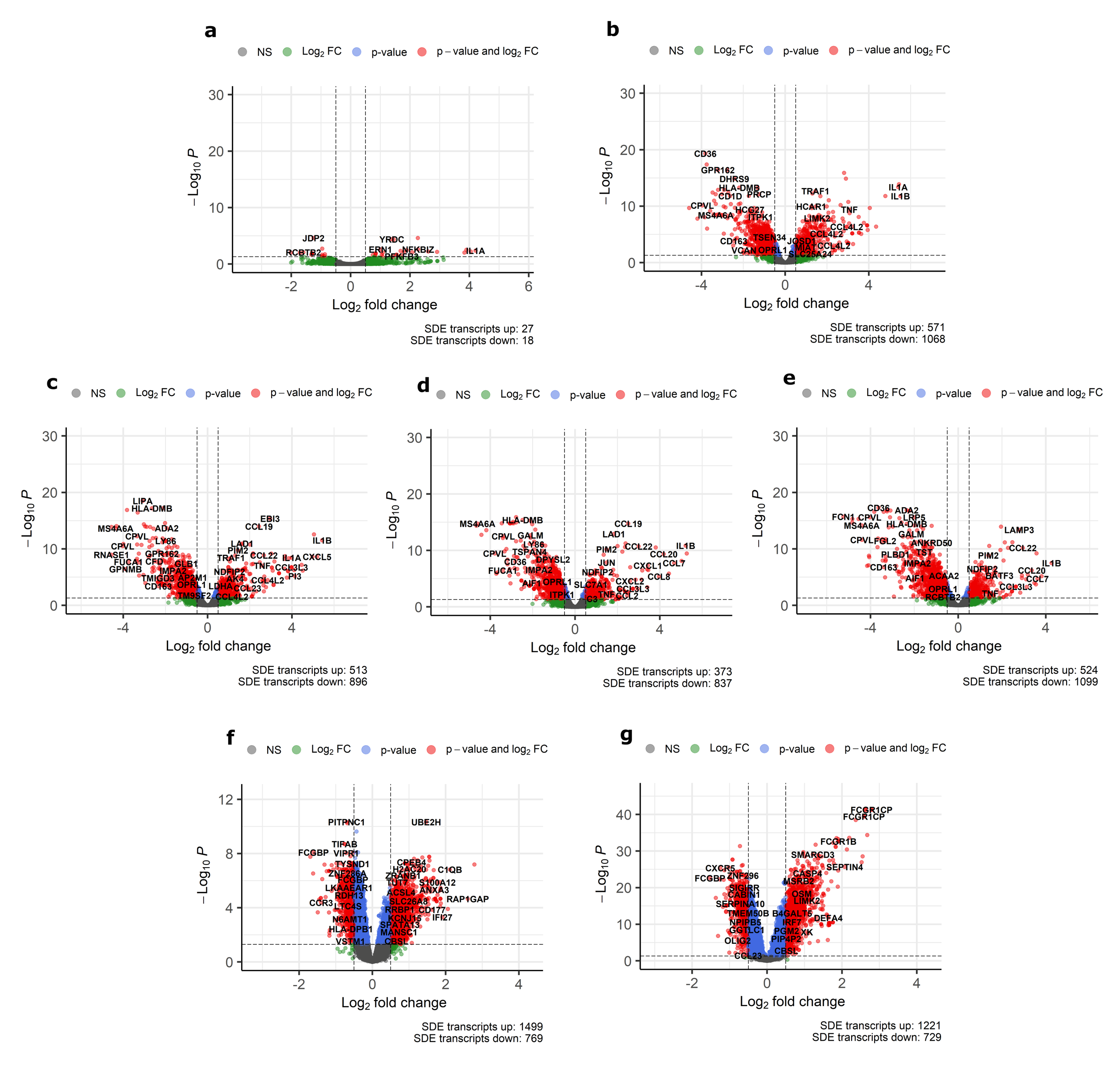

**Supplementary Figure 2: Differential transcript expression**

Volcano plots showing differential transcript expression in the *in vitro* WBA infection model between *Mtb* infection and controls (uninfected) in the data sets used for the analysis.
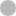
 - not significant,
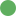
 - log_2_FC > 0.5 or < ‒ 0.5,
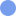
 - p-value < 0.5,
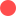
 - both p-value and log_2_FC significant. Differential expression at a) 6 h, b) 24 h, c) 48 h, d) 72 h and e) 96 h post-*Mtb*-infection *in vitro*, compared to the respective uninfected control. Differential expression of *in vivo* TB disease in f) paediatric and g) adult patients, compared to LTBI control patients.

**Supplementary Figure 3: Disco plots for 96 h concordance comparisons**

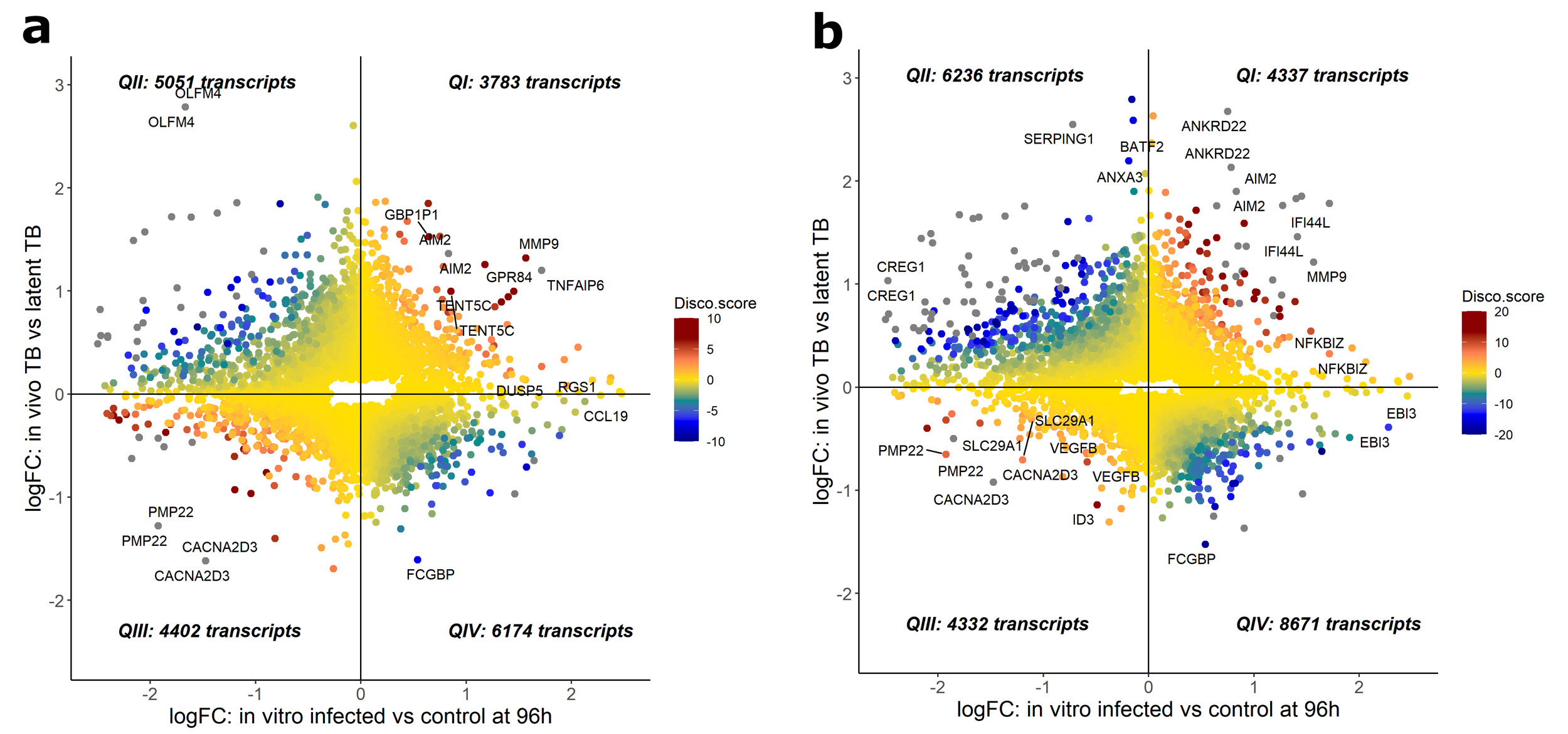

Evaluation of concordance of gene expression changes in a) adult TB and *in vitro* 96 h and b) paediatric TB and *in vitro* 96 h. Disco.score shown by colour intensity, ranging from blue (low concordance) to dark red (high concordance). Transcripts found in each disco plot are segmented into four quadrants (Q) as follows; QI - concordantly upregulated, QII - discordant, upregulated *in vivo* & downregulated *in vitro*, QIII - concordantly downregulated and QIV - discordant, downregulated *in vivo* and upregulated *in vitro*. Numbers of identified transcripts are shown in each quadrant.

**Supplementary Files:**

GO_pathways_in_vitro.xlsx

GO_pathways_in_vivo_adult.xlsx

GO_pathways_in_vivo_paediatric.xlsx
